# Accelerating Biomolecular Modeling with AtomWorks and RF3

**DOI:** 10.1101/2025.08.14.670328

**Authors:** Nathaniel Corley, Simon Mathis, Rohith Krishna, Magnus S. Bauer, Tuscan R. Thompson, Woody Ahern, Maxwell W. Kazman, Rafael I. Brent, Kieran Didi, Andrew Kubaney, Lilian McHugh, Arnav Nagle, Andrew Favor, Meghana Kshirsagar, Pascal Sturmfels, Yanjing Li, Jasper Butcher, Bo Qiang, Lars L. Schaaf, Raktim Mitra, Katelyn Campbell, Odin Zhang, Roni Weissman, Ian R. Humphreys, Qian Cong, Jonathan Funk, Shreyash Sonthalia, Pietro Liò, David Baker, Frank DiMaio

## Abstract

Deep learning methods trained on protein structure databases have revolutionized biomolecular structure prediction, but developing and training new models remains a considerable challenge. To facilitate the development of new models, we present AtomWorks: a broadly applicable data framework for developing state-of-the-art biomolecular foundation models spanning diverse tasks, including structure prediction, generative protein design, and fixed backbone sequence design. We use AtomWorks to train RosettaFold-3 (RF3), a structure prediction network capable of predicting arbitrary biomolecular complexes with an improved treatment of chirality that narrows the performance gap between closed-source AlphaFold3 (AF3) and existing open-source implementations. We expect that AtomWorks will accelerate the next generation of open-source biomolecular machine learning models and that RF3 will be broadly useful as a structure prediction tool. To this end, we release the AtomWorks framework (https://github.com/RosettaCommons/atomworks), together with curated training data, code and model weights for RF3 (https://github.com/RosettaCommons/modelforge) under a permissive BSD license.

## 1 Main

Deep learning has revolutionized the prediction of single chain proteins and their interactions, extending recently to include a wider set of biomolecules including DNA, RNA, and small molecules[1, 2, 3, 4, 5, 6, 7]. However, open-source efforts to develop and refine biomolecular machine learning models have been hamstrung by challenges that hinder progress. First, the heterogeneous data quality, annotations, and sources impose a considerable engineering overhead. Prospective researchers must spend months preparing data, optimizing processing, and battling edge cases within the Protein Data Bank (PDB)[8] before training any networks or, more commonly, opt to only handle a subset of the field’s complexity (e.g., protein-only models [9, 10, 11]). Second, the reliance on bespoke data pipelines for each network prevents generalization across use cases. Operations developed for one network cannot easily transfer to another, leading to duplication of code and effort across the research community [12, 13].

We reasoned that a modular, research-centric framework that enables rapid prototyping of protein structure prediction and design models would accelerate progress. We set out to develop such a framework within a high-performance codebase and utilize it to train open-source structure prediction and design methods.

## 2 Democratizing foundation modeling on biomolecular structure data

We describe below the tenets that guide development of AtomWorks, our generalized computational framework for biomolecular modeling. AtomWorks enables rapid prototyping and scalable training of biomolecular foundation models within one unified framework, emphasizing high-quality data handling, comprehensive documentation, and extensive testing to democratize access to biomolecular foundation modeling.

### AtomWorks prioritizes high-quality data

Data quality is a key determinant of machine learning model performance. Within structural biology, the PDB is the primary experimental dataset, supplemented increasingly by model-derived distillation datasets as well [1, 5, 9]. These data sources, however, contain numerous edge cases that arise from the inherent heterogeneity of structural biology data. Correctly identifying and resolving these edge cases often requires expertise in biochemistry and structural biology. In AtomWorks, we begin by standardizing inputs with proper handling of these myriad data challenges; for example, identifying and removing leaving groups, correcting bond order after nucleophilic addition, fixing charges, parsing covalent geometries, imputing missing coordinates, and appropriate treatment of structures with multiple occupancies and ligands at symmetry centers. We find that our input processing approach translates to higher-quality derived downstream features. For example, AtomWorks reference conformers – generated through an empirical force field energy-minimization that is sensitive to erroneous charge and bond annotations – have lower energies than the reference conformers of another fully open-source model [7](fig. S2; median 67 vs 109 kJ/mol, as calculated by PoseBusters software suite [14]).

### AtomWorks maximizes research velocity by enabling rapid prototyping

Current biomolecular modeling networks rely on independent data loading and featurization pipelines despite training procedures with many shared operations. To reduce duplicated efforts, we split subsequent data processing and featurization into modular components that operate on our common atom-level representation of structure. We further separate our logic into general-purpose operations that derive common features (e.g., load multiple sequence alignments, identify symmetric chains, generate reference conformers with chemoinformatics software such as RDKit [15]) and operations that translate the derived features into model-specific tensors. We find this approach both facilitates reuse of core building blocks across networks and reduces the complexity of adding new features (fig. S8). Previously, pipelines were single functions that incrementally converted input features into model tensors, discarding unneeded information along the way. Adding new features to such models involved introducing significant complexity to the code, leading to unmanageable data processing pipelines. Within the AtomWorks paradigm, the output of each step is not an opaque dictionary with model-specific tensors but instead an updated version of our atom-level structural representation (built on the open-source Biotite library’s AtomArray [16]). Operations within – and between – pipelines thus maintain a common vocabulary of inputs and outputs. Researchers need not contend with all existing features; they can focus on their specific experimental hypotheses, with knowledge that any modifications to the internal state (e.g., re-ordering or deleting atoms) will be appropriately handled downstream.

### AtomWorks enables the scalable training of biomolecular prediction and design models

Building on this foundation, we implemented AtomWorks pipelines to train RF3, RF All-Atom (RFAA) [4], LigandMPNN [17], ProteinMPNN [18], and a to-be-published all-atom generative model (fig. 1). We find that the AtomWorks framework allows most code (>80%) to be shared across networks; researchers can repurpose existing components to rapidly test hypotheses, and improvements to common operations simultaneously benefit all methods. We contrast this approach to the prior paradigm where each model existed within a siloed codebase; component sharing was accomplished by copy-and-paste followed by modification to meet pipeline-specific needs. Indeed, the modular architecture of AtomWorks allows replacing 2,000+ lines of code in LigandMPNN with a 100-line declarative pipeline that borrows from operations originally written for RF3 and RFAA (fig. 1). Critically, our framework is also highly efficient. By relying on vectorized C operations and the pre-optimized Biotite library, we can parse from source structural files in milliseconds. This efficiency allows us to, for example, process a 6,000 token batch through our LigandMPNN pipeline within the time of a single forward/backward pass (0.6s).

**Fig. 1:**
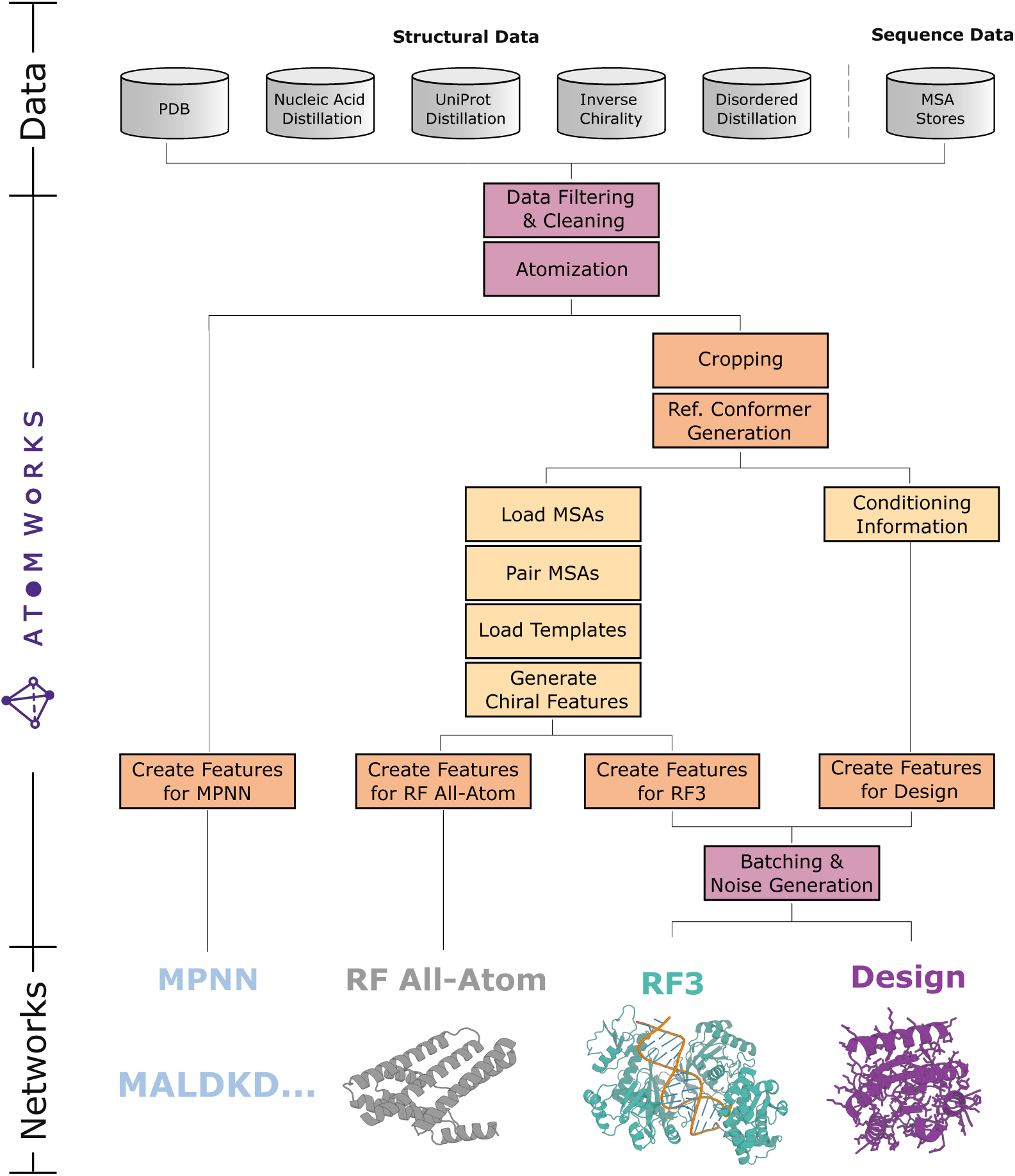
AtomWorks enables training of state-of-the-art models for structure prediction, generative protein design, and fixed-backbone sequence design within a single data framework. **Top**: AtomWorks can ingest (and RF3 is trained on) a diverse set of datasets including the Protein Data Bank (PDB), generated nucleic acid distillation sets, monomer distillation sets, PDB structures with inverted chirality, and PDB structures with extended disordered regions. **Middle**: Networks for all biomolecular modeling use cases can share common components within the AtomWorks framework. **Bottom**: Representations of outputs from various models trained with AtomWorks.

### AtomWorks follows industry-grade testing and documentation practices

We developed AtomWorks not as a point-in-time artifact but instead as a living library to be extended and improved. Currently, expert knowledge regarding how to handle the full complexity of biomolecular data is concentrated within a handful of academic labs and commercial entities. To address this disparity, we release alongside AtomWorks industry-grade tests for all operations (>85% coverage) and comprehensive documentation with worked examples illustrating how to develop pipelines for biomolecular modeling tasks. By focusing on usability, we facilitate the transfer of knowledge so that future researchers can more easily benefit from our foundation.

## 3 Training RF3

We use the AtomWorks framework to train RF3, an all-atom biomolecular structure prediction network competitive with leading open-source structure prediction networks. By including additional features at train-time – implicit chirality representations and atom-level geometric conditioning – we improve performance on tasks such as prediction of chiral ligands and fixed-backbone or fixed-conformer docking.

### RF3 simplifies dataset integration via AtomWorks

As AtomWorks supports loading directly from raw crystallographic information files (CIF) through a unified processing pipeline, all that is needed to include additional training datasets are directories of predicted structures and, optionally, the corresponding MSAs. For RF3, we leverage this flexibility to introduce a number of novel distillation datasets. First, we incorporate a previously-reported distillation set of multi-domain proteins from AFDB truncated into pairs of interacting domains that mimic protein-protein interfaces [19, 20]. We also develop two new nucleic acid distillation datasets: a protein-nucleic acid complex distillation set and an RNA distillation set (with 27K examples and 10K examples, respectively; see supplemental methods) (fig. S1). Further, to address the issue of hallucinated secondary structure, we introduce a disordered distillation set that uses the Rosetta macromolecular modeling software [21] to generate structures with “extended” backbones for disordered regions (fig. 3b).

### RF3 accurately adheres to specified stereochemistry out-of-the-box, without inference-time guidance

To resolve the well-documented issue of incorrect chirality within diffusion-based structure prediction models, we represent stereochemistry by the sign of the angles formed by the atoms surrounding each chiral center. At each denoising step, we provide the diffusion model with a vector feature of the gradient of the error of the ideal angle (fig. S7, [4]). In addition, to encourage the model to respect the chiral features, we invert the chirality in 2% of PDB examples as a train-time data augmentation. We find that after making these improvements, the network predicts the correct chirality for 88% of ligand chiral centers within our test set, compared to 84% for AF3 and 76% for Boltz-2 without inference-time guidance (fig. 2a).

**Fig. 2:**
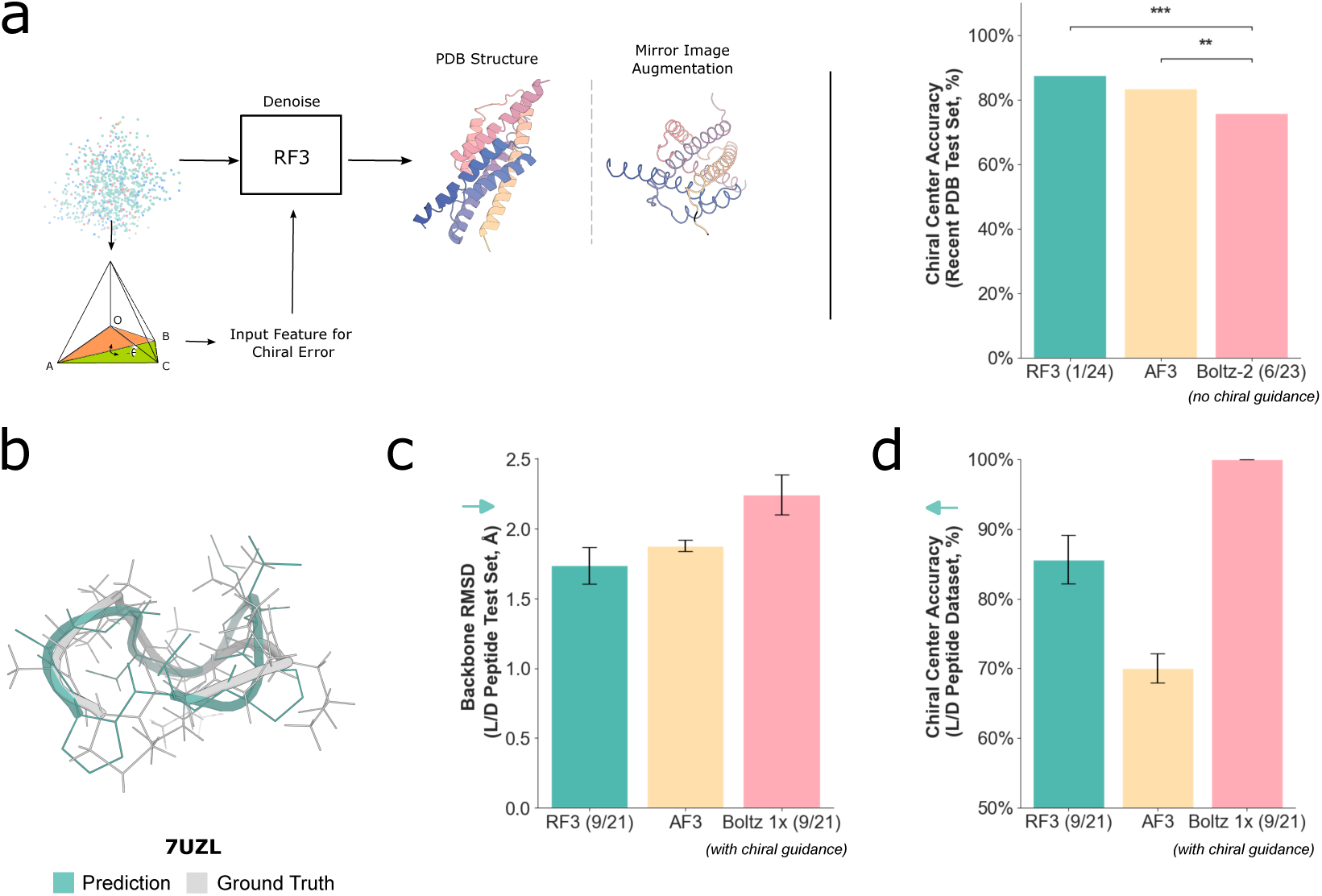
RF3 respects the chirality of the inputs. **A**: RF3 respects the chirality of small molecule inputs more than all other structure prediction models. We first cluster all small molecules in our test set by CCD code and average the percentage of correctly predicted centers within each cluster. Overall chiral center accuracy is computed by taking the mean across all clusters. Statistical significance was assessed using two-sample t-tests. RF3 and AF3 both significantly outperformed Boltz-2 (p < 0.001 and p = 0.002, respectively), while the difference between RF3 and AF3 was not significant (p = 0.064). Model training date cutoff indicated in parentheses. **B**: Comparison of predicted (teal) with ground-truth crystal structure (gray) for PDB ID 7UZL. In this example, 100% of the chiral centers are predicted with the correct chirality (including three D amino acids). **C**: On a test set of mixed chirality macrocycles from 2022, outside the training date cutoff of all models benchmarked, RF3 predicts the structures with a high degree of accuracy (1.74 mean backbone RMSD) **D**: In the mixed chirality test set, 85% of chiral centers are predicted correctly by RF3 vs. 70% by AF3. Boltz 1x structures were predicted with inference-time chiral guidance and thus are guaranteed to satisfy the input chirality.

**Fig. 3:**
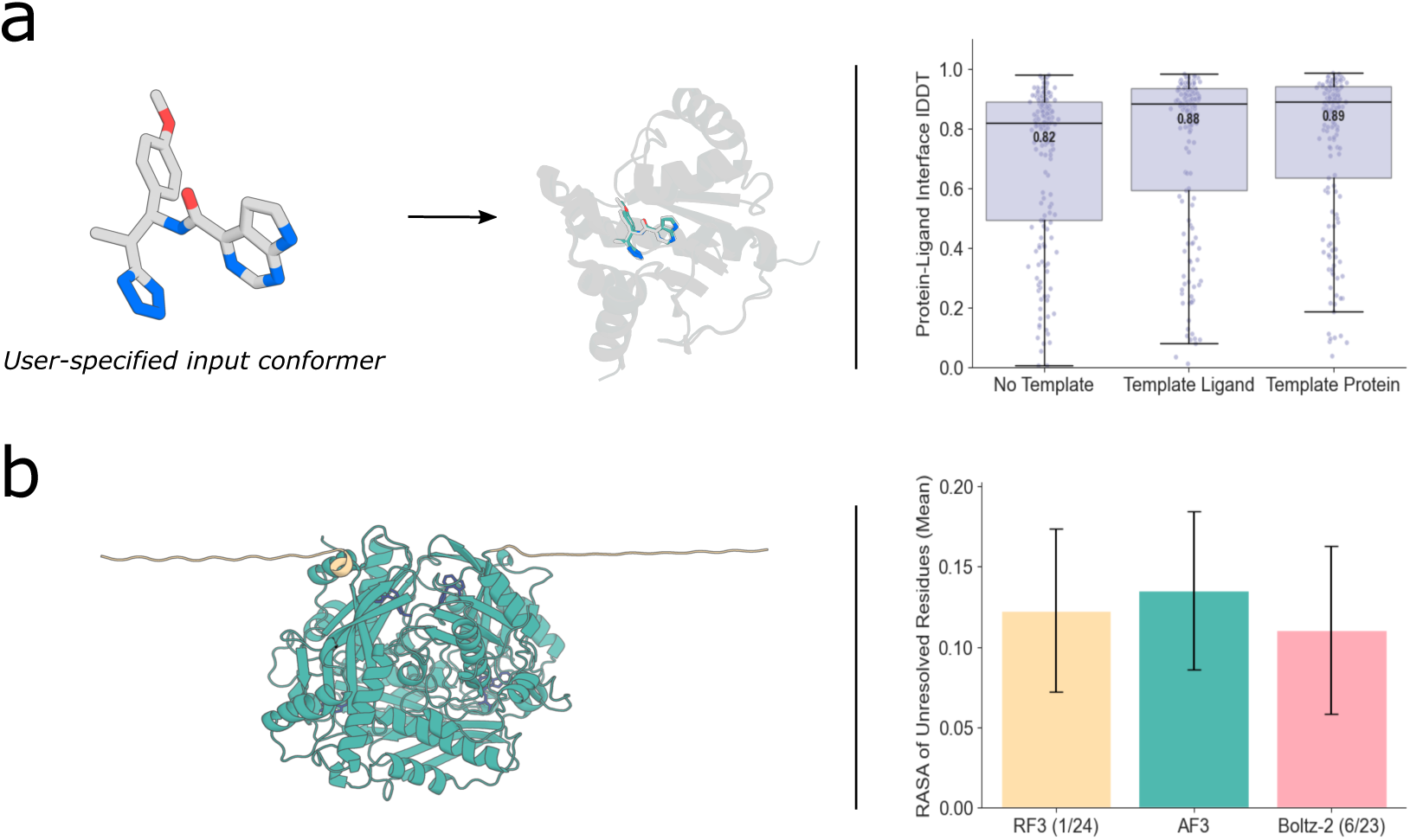
Novel capabilities of RF3. **A**. RF3 enables users to input desired conformers. (left) example of a user input conformer and prediction. (right) providing the ligand ground truth ligand conformer or the protein holo conformation improves accuracy. “Templating” in this case means providing all-by-all pairwise distances within the ligand or protein. **B**. RF3 is trained with a disorder distillation set. In contrast to AF3, which repredicted the entire PDB with AF2 to show examples of extended disordered regions, we chose to use the more compute-efficient Rosetta macromolecular modeling software to generate structures with “extended” backbones for disordered regions. In 2% of cases, the model is trained with PDB examples with extended disordered regions. (left) example of a prediction with a large unresolved region which is predicted in an extended conformation. (right) Analysis of the mean RASA over unresolved regions in our test set. Model training date cutoff indicated in parentheses.

**Fig. 4:**
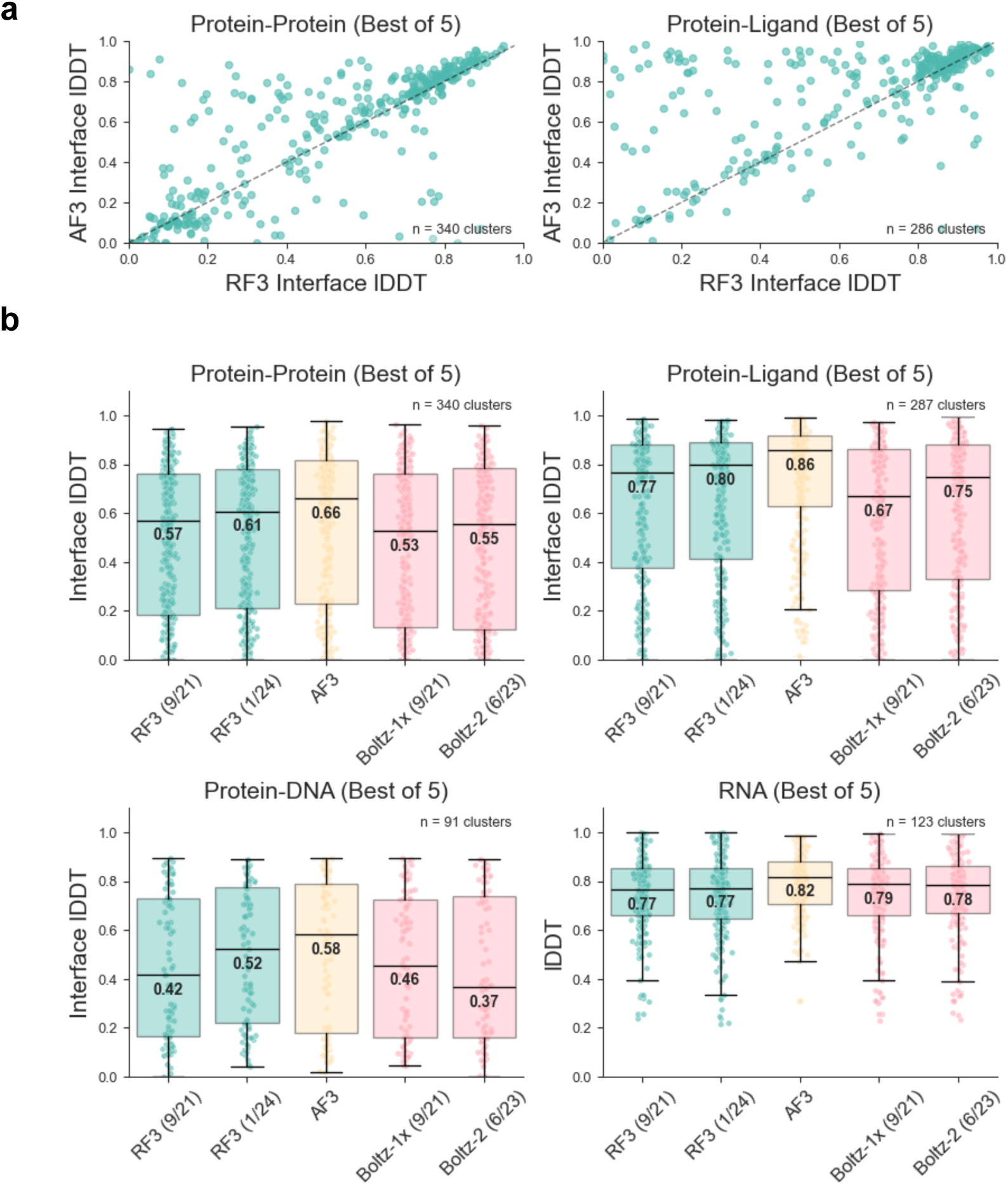
RF3 accurately predicts biomolecular interactions. **A**. Scatterplots comparing all-atom interface lDDT of RF3 and AF3 on protein-protein interactions and protein-ligand interactions. To reduce redundancy, the test dataset was clustered (by sequence homology 40% for polymers and CCD identity for non-polymers); each point represents a cluster mean. For all networks, we generate five structures from the same seed and take the sample that scores the highest on each metric (“Best of 5” approach). **B**. Boxplots showing accuracy of RF3, AF3 and Boltz on different structure modeling tasks. Two versions of RF3 are shown: one trained on structures released before September, 2021 and another trained on structures deposited before January, 2024. Each point in the boxplot is a mean value over a cluster of structures. Model training date cutoff indicated in parentheses.

We further reasoned that the improved handling of chirality would enable RF3 to accurately predict mixed L/D peptides, a growing therapeutic class of molecules noted for their proteolytic stability and structural diversity. On a set of cyclic mixed L/D peptides from after the training date cutoff of all methods [22], we find that RF3 predicts 86% of chiral centers correctly (compared to 70% for AF3 and 100% for Boltz-1x, when using inference time guidance to enforce chirality; 2d). Moreover, the structures are highly accurate (mean 1.74Å backbone RMSD vs. 1.88 for AF3 and 2.24 for Boltz-1x; 2c). We hypothesize that inference-time modifications may shift the network outside the training distribution, which would explain why we find improved accuracy from representing chirality as a learned feature.

### RF3 enables flexible user control through arbitrary atom-level conditioning

Users may specify distances between atoms that they want recapitulated in the output structure to enable incorporation of experimentally derived constraints, protein- or ligand-docking against a known protein structure, or protein folding around a specific ligand conformer. To measure the ability of the network to adhere to these constraints, we test the network on *holo* structure docking and protein folding around a rigid conformation of a ligand. When folding the protein around a rigid ligand, median protein-ligand interface accuracy increases from 0.821 to 0.882. Similarly, when providing distances constraining the *holo* structure, protein-ligand interface accuracy improves from 0.821 to 0.890 (fig. 2a). In all cases of templating the ligand, the model respects the given small molecule conformation (median ligand-only lDDT 0.991, fig. S6).

### RF3 narrows the performance gap between existing open-source structure prediction and AF3

Despite recent advances, open-source models continue to trail AF3 performance on most biologically relevant problems. In fig. 4, we compare the performance of RF3 to AF3 and another open source prediction network, Boltz [7]. All methods were run with a single seed, inference parameters to generate 5 diffusion predictions from one run of the trunk, and identical MSAs; for stability, we select the best-scoring sample for each metric (choosing a single structure with the confidence head shows similar trends, fig. S4). We find that the performance of RF3 is between the closed-source AF3 and open-source Boltz in almost all categories. When the training date cutoff is extended to 1/2024, we find RF3 performance increases across-the-board (0.571 vs. 0.607 median protein-protein interface lDDT, 0.766 vs 0.798 median protein-ligand interface lDDT, 0.415 vs 0.523 median protein-DNA interface lDDT and 0.765 vs. 0.772 median RNA only lDDT).

Antibody-antigen complex prediction represents a specific structure prediction use case with widespread applications within the pharmaceutical industry. When evaluated on a clustered, de-leaked test set of antibody-antigen complexes from the recent PDB, RF3 again performs between AF3 and the other methods (in this case, Boltz-2 and Chai-1): 33% of examples achieve DockQ > 0.23 for RF3, compared to 44% for AF3, 22% for Boltz-2, and 28% for Chai-1 (fig. S3).

## 4 Discussion

AtomWorks lowers the barrier-to-entry for training machine learning models on data from the Protein Data Bank and other structural biology databases, and RF3 sets a new standard for open-source models for prediction of the structures of protein-protein interfaces, protein-ligand interactions, and mixed L/D peptides. Using the AtomWorks modular data pipeline, scientists can develop new deep-learning networks without extensive software development experience. Our improved data processing directly translates to better generalization — when subset to cases with fewer than five similar ligands in the PDB, we find that reference conformer energies correlate with prediction accuracy between RF3 and Boltz (fig. S2). At the Institute for Protein Design, AtomWorks and RF3 have expanded both the number of developers who can contribute to deep learning projects and the code sharing between prediction and design efforts. We anticipate that these packages will be widely useful for new model development and to that end we release all data, code, and model weights for open use by the community.

## Acknowledgments

We thank L. Goldschmidt and K. VanWormer for maintaining the computational and wet lab resources at the Institute for Protein Design. We thank Microsoft for the generous donation of Azure compute credits which enabled training all of the networks described in this work. We also thank Minjia Zhang, Xinyu Lian, and Hoa La who helped with GPU acceleration for early versions of RF3, Jason Yim for helpful discussion on generative modeling and Roy Tal-Dew, Ben Fauber and Hari Sadasivan for advice on acceleration via cuEquivariance.

## Author Contributions

N.C., R.K, and S.M. co-led the effort and may change the order of their names for personal pursuits to best suit their own interests. Research design: R.K., N.C., S.M., F.D., D.B.; Development of AtomWorks library: N.C., S.M., R.K., Ma.K., R.B., F.D., K.D., R.M., L.M., B.Q., R.W., J.F., A.N., P.S.; Development of RF3: F.D., R.K., W.A., N.C., T.T., S.M.; Evaluation of RF3 on different structure prediction tasks: M.B., N.C., F.D., R.B., O.Z., K.C., M.K.; Development of distillation datasets: F.D., L.M., A.F., Me.K., I.H.; Contribution of code and ideas: M.B., J.B., Y.L., L.S., Ma.K., A.N., S.S.; Development of LigandMPNN integrated with AtomWorks: A.K., N.C..; Supervision throughout the project: D.B., F.D., R.K.; Writing - original draft and figures: N.C., R.K., S.M., K.D., Y.L., L.S., F.D., D.B.; Writing - review and editing: All authors.

## A Supplement

### A.1 Naming Conventions

We adopt a consistent, composable naming convention for different “bits” of a `mmCIF` file throughout data parsing, preprocessing, loading, and featurization so that our code remains unambiguous. We outline the conventions below; they are also described within the AtomWorks documentation.

#### Entities vs. Instances

Within our nomenclature, *entities* are chemical compounds where we distinguish the (covalent) connectivity and components, but not the coordinates. *Instances*, meanwhile, are unique copies of an entity in 3D. In Python terms: *entity* ∼ *class* and *instance* ∼ *instance of that class*.

For example, within a `mmCIF` file, there may be multiple copies of the same chain (sometimes referred to as `asym_id` in PDB files), each with a unique set of coordinates, but identical sequences and connectivities. These compounds are distinct *instances*, but the same underlying *entity* (e.g., same `UNIREF` identifier).

#### Suffixes

- _`entity`: A unique numeric id for each entity.
- _`id`: A group id that may or may not correspond to more than one instance, subdivided, for example, through symmetries during assembly building. For example, we would consider the PDB’s `asym_id`: to be an _`id`, as it uniquely specifies the entity, but not the instance (due to transformations). If unfamiliar with transformations and biological assemblies in the PDB, consult the RCSB documentation before continuing.
- _`iid`: The “instance ID,” which uniquely specifies a group of atoms in three-dimensional space.

#### Chains, PN_Units, and Molecules

- **Chain.** The smallest covalently bound unit within the PDB is the “chain,” with each chain represented in a `mmCIF` file by a unique combination of an `asym_id` and a transformation_id.
- **PN_Unit.** Short for “polymer or non-polymer unit.” We define a `pn_unit` as covalently linked chains of the same type. For example, an oligosaccharide may be represented as multiple non-polymer chains covalently bound together, which we should treat as one `pn_unit`. However, an oligosaccharide bound to a protein would be two separate pn_-units (one for the oligosaccharide, one for the protein), as they differ in chain type.
- **Molecule.** Aligned with the definition of a molecule in chemistry (created by traversal of the bond graph). It refers to a single connected component of a covalent bond graph. May contain multiple `pn_units` (e.g., a covalent modification of a protein with a glycan would be 2 `pn_units` but 1 molecule).

### A.2 Datasets

#### A.2.1 Dataset Preprocessing

For our PDB dataset, we follow [5] and pre-process the full PDB into a dataset that represents monomers and a dataset that represents interfaces. In our case, we call the monomers dataset “`pn_units`” and the interfaces dataset “interfaces.“

Namely, we:

- Resolve clashes (defined as a pair of chains with any heavy atoms within 1 Å) by removing the smaller chain
- Remove chains with misannotated bonds (e.g., oxygen-oxygen bonds, flourine-flourine bonds, bonds involving free oxygen or hydroxyl groups)
- Remove fully unresolved chains
- Collate additional metadata from the CIF file (e.g., subject of investigation labels, resolution, EC class, ligand fit-to-density, etc.)

After running this pre-processing pipeline, we are left with two metadata dataframes - one of all the chains (more precisely, `pn_units`) in the PDB, and one with all the interfaces in the PDB. We sample from these two dataframes during training; we also find that these dataframes are generally useful for querying and analyzing the PDB outside of a model training context. We release both dataframes, alongside the corresponding preprocessing code, within AtomWorks.

#### A.2.2 Distillation Datasets

We introduce two novel distillation datasets: an RNA Distillation dataset and a Nucleic Acid Complex distillation dataset (fig. S1). We also use previously published distillations sets of AF2 predicted monomers [23] and domain-domain interactions that was collated from the AlphaFoldDB [20, 19].

##### RNA Distillation

An initial set of RNA sequences was obtained from RNAcentral (January 2025 release) [24], restricted to sequences of length 90-500 nucleotides. To enrich for structured noncoding RNAs, sequences were filtered using EternaFold [25] secondary structure predictions, in order to retain only sequences with ≥ 70% of positions predicted to be base-paired. This set was clustered using MMseqs2 (90% sequence identity, 80% coverage) [26]. From each cluster, a single representative sequence was selected based on the highest predicted base-pairing fraction. Structures were predicted using an earlier version of RF3, grouped into bins based on length (5-nt bins), and filtered by output confidence metrics. Predictions were selected based on the union of two criteria: (1) the five predictions with the lowest predicted design error (pDE) scores per length bin, and (2) all predictions with pDE scores below a length-adjusted threshold defined as pDE ≤ ln(0.075 · length). A final filter was applied to retain only sequences with pLDDT ≥ 0.62 and pAE ≤ 15.

##### Nucleic Acid Complex Distillation

Protein sequences and cognate DNA sequences were obtained from multiple sources, including cis-BP [27], UniProbe[28, 29], TRANSFAC [30], and multiple HT-SELEX and ChIP-Seq studies. Proteins were grouped by family using a combination of provided annotations and sequence alignment / clustering. DNA-binding domains were manually identified for representatives of each protein family using a combination of source annotations, UniProt domain annotations, and visual inspection of structures from the AlphaFold Protein Structure Database; domains for the rest of the examples were determined by alignment to these representatives. DNA sequences were chosen from the sequences tested experimentally in the raw data, either the top-ranked in the original experiment or the best matches to the consensus motif as determined by MEME. Structures of the protein-DNA complexes were predicted using either RFNA [31], a fine-tuned version of RFallatom [4], or RF-3, then filtered for high-confidence (mean pLDDT > 80, mean interface PAE < 10).

#### A.2.3 Implemented Transforms

Within adjacent fields such as computer vision and language modeling, researchers have developed libraries of `Transforms` for common data loading operations. For example, the popular Torchvision library contains common operations such as scaling, normalization, and cropping of images prior to converting inputs into PyTorch tensors. Such a framework accelerates research velocity, as `Transforms` can be chained together to rapidly test new ideas.

Inspired by the success these `Transform`-based approaches, we developed a Transform framework within AtomWorks tailor-made for biomolecular structures. Rather than relying on pixels as a common representation, we make use of the `AtomArray` — an annotated list of atoms implemented in C-level vector operations within the open-source Biotite library [16]. `Transforms` thus operate on `AtomArrays` — they receive as input an `AtomArray` and return a modified `AtomArray`. We find that this approach immensely simplifies the addition of new features while also enhancing the readability of our code (fig. S8).

A full description of our library is available within the AtomWorks documentation, along with worked examples.

### A.3 Architecture Details

#### A.3.1 Model Inputs

Inputs to RF3 include the primary polymer sequence, evolutionary information (both protein and RNA MSAs), atom-level ground-truth templates (a generalization of AF2-style protein templates), reference conformers generated with RDKit (one per residue), chiral features (described in A.3.3), and ML force field embeddings (optional) (fig. S1). For training, we compute MSAs with a combination of HHBlits [32] for the pre-2021 MSAs and MMSeqs2-GPU for the post-2021 MSAs [33]. We did not directly ablate the impact on training of using MSAs from HHBlits vs. MMSeqs2GPU.

For the ground-truth templates, we embed information into the model in two ways: a) via the token-level template track, binning pairwise distances into 64 distance bins, following [4]; and, b) via the reference conformer, adding an additional atom-level feature that is directly embedded within the `AtomAttentionEncoderDiffusion`. At inference time, either approach could be used; they each have distinct advantages. The results shown in fig. 3 and fig. S6 employ both form of templating to maximally constrain geometries.

It may be possible to leverage progress in the ML force literature — where recent models train on millions of small molecules and achieve performance approaching traditional quantum mechanical methods — to enhance performance on protein small-molecule complexes [34]. As a preliminary test, we added learned embeddings of small molecules from a MLFF based on the MACE architecture (Egret-1 [35]) to the network as atom-level features during fine-tuning. In particular, we sample 8 conformers per residue, embed these conformers with Egret-1, and then feed the embeddings to our `ConformerWeightedAverage` block (algorithm 2). We find our approach did not significantly improve accuracy over the standard method of directly embedding conformer coordinates and inverse distances (table S1). However, we believe this area merits further exploration.

**Table S1:** Best-of-1 local distance difference test (lDDT) performance by molecular category. Comparison of Egret embeddings (With MACE) and Baseline (Without MACE), best-of-1 (random choice; not confidence).

| Metric (IDDT) | Best-of-1 With MACE (Egret) | Best-of-1 Baseline |
| --- | --- | --- |
| RNA | 0.719 | 0.724 |
| Ligand | 0.919 | 0.917 |
| Protein | 0.821 | 0.822 |
| Protein-DNA | 0.392 | 0.404 |
| Protein-Ligand | 0.685 | 0.684 |
| Protein-Protein | 0.465 | 0.466 |

#### A.3.2 Improving training stability

We began by implementing the algorithms described in the Supplemental Methods of the AlphaFold3 (AF3) manuscript [5]. All the training details were not described in the AF3 paper and we anticipate that our training procedure has some discrepancies from those used in the AF3 weights that were released for academics by Deepmind. During this process, we identified two modifications that were essential for achieving high training accuracy.

First, we replaced the loss weighting term used in the AF3 supplement,

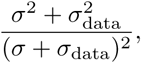

with a more faithful implementation of the EDM loss weighting from [36]:

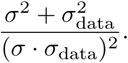

Second, we updated the residual connection in the diffusion transformer. The AF3 supplement specifies the residual connection between operations in the diffusion transformer as:

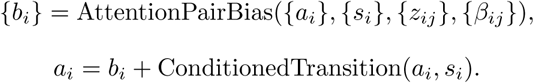

We replaced this with a more modern transformer residual connection:

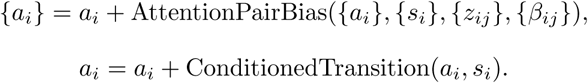

When training with this residual connection, we found that in some early experiments the magnitude of some of the weights before the attention operation started to get larger than expected causing the attention weights to look one-hot encoded after the Softmax operator. We noticed a similar result in [37], and adopted their solution - applying a LayerNorm to the queries and keys before the attention operation. We also note that this solution (known as QK-norm) has become pervasive in the LLM literature as well.

#### A.3.3 Chirality Features

We largely follow the chiral featurization of [4] (similar models of chirality are reported in [13, 38]). Chiral centers are often categorized using a binary label of (r) and (s) which is computed by assessing a “priority” on each atom in the tetrahedron. The lowest priority atom is pointed away from the user and the remaining three atoms are ordered by decreasing priority. For cases where the priority decreases in a clockwise direction, the stereocenter is assigned (r) and for counter-clockwise direction, the stereocenter is assigned (s).

We felt the model would have difficulty learning the arbitrary priority list governing (r) and (s) assignments. Therefore, we opted to present the chirality to the network *geometrically*. For each non-H atom bonded to the chiral center, we compute a vector representing the difference between the current atom position, and where that atom should go given the positions of other atoms comprising the chiral center. Formally, we use pseudotorsions to define the chiral center. The pseudotorsion specifies the planarity of the fourth atom and we use that feature to drive the structure towards a positive angle or negative angle depending on the input stereochemistry.

We take advantage of this property of dihedral angles to specify chirality to the network. The dihedral angle between planes (*v*_1_, *v*_2_, *v*_3_) and (*v*_2_, *v*_3_, *v*_4_) will be positive in the first case and negative in the second. In practice in RF3, we do not always have access to all four substituents of a chiral center (one of them could be a hydrogen which is not explicitly modelled). Despite this we do have sufficient information to determine the chirality of a given system given three points since we know the chiral center has coordinates: *o* = (0, 0, 0). We can then construct planes consisting of (*o, v*_1_, *v*_2_) and (*v*_1_, *v*_2_, *v*_3_), and compute the dihedral angle between them which will be either arcsin 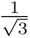 or − arcsin 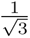 (0.6155 radians or −0.6155 radians).

To provide the ideal reference angle as a feature to the network, we compute the analytical gradient of the error of the noisy structure atom dihedral angle to the reference angle with respect to the coordinates. Formally,

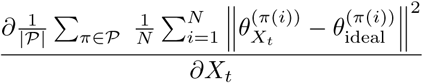

where *P* is the set of all permutations of explicitly represented atoms with the chiral center as the first point in the dihedral computation, N is the number of chiral centers, *X_t_*is the noisy structure going into the network, and *θ_Xt_* is the computed dihedral based on the noisy structure.

In practice, we replaced the AtomAttentionEncoder in the Diffusion Module with the following algorithm. We leave the AtomAttentionEncoder block in the pairformer untouched.

##### Algorithm 1 AtomAttentionEncoderDiffusion

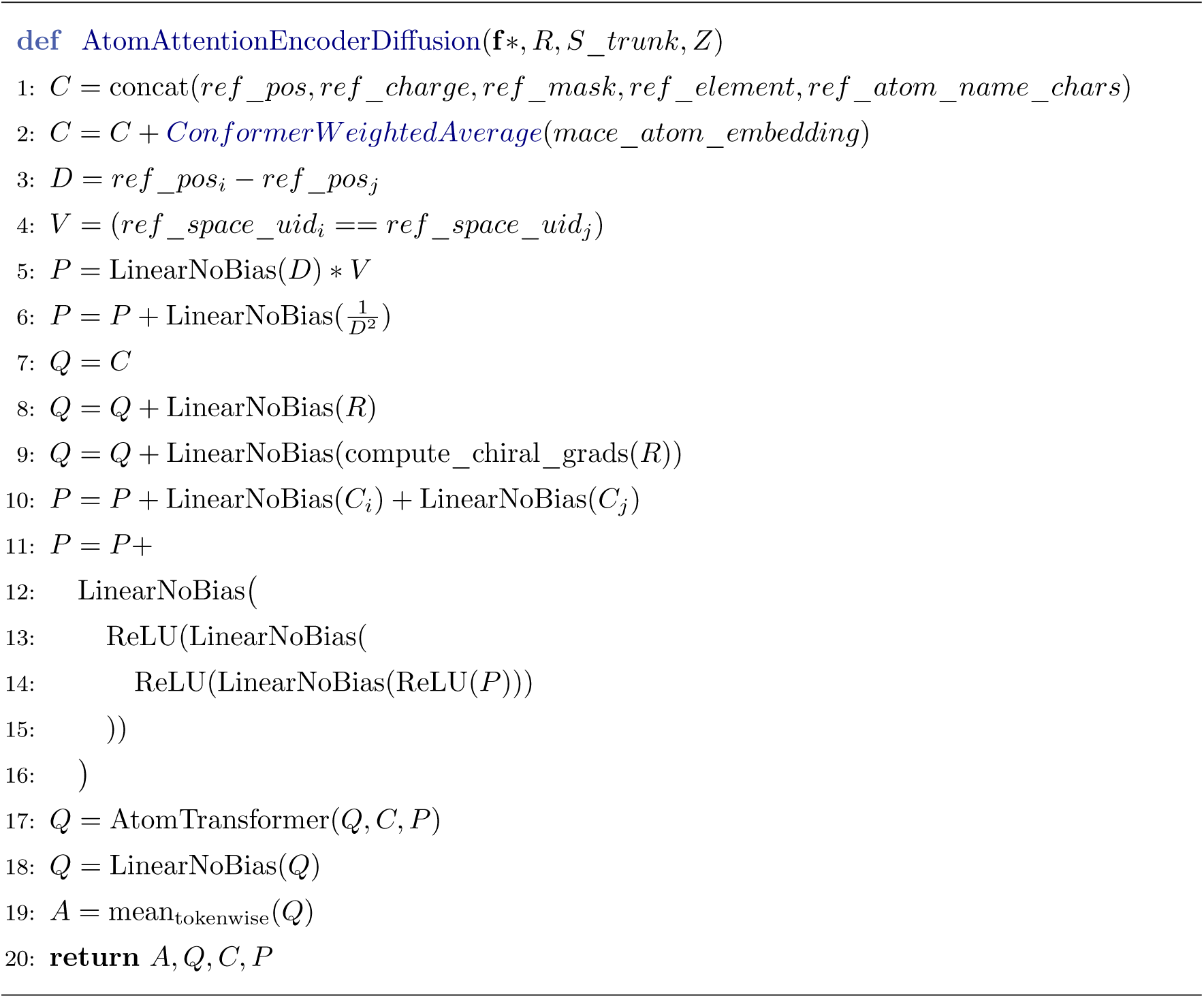

##### Algorithm 2 ConformerWeightedAverage

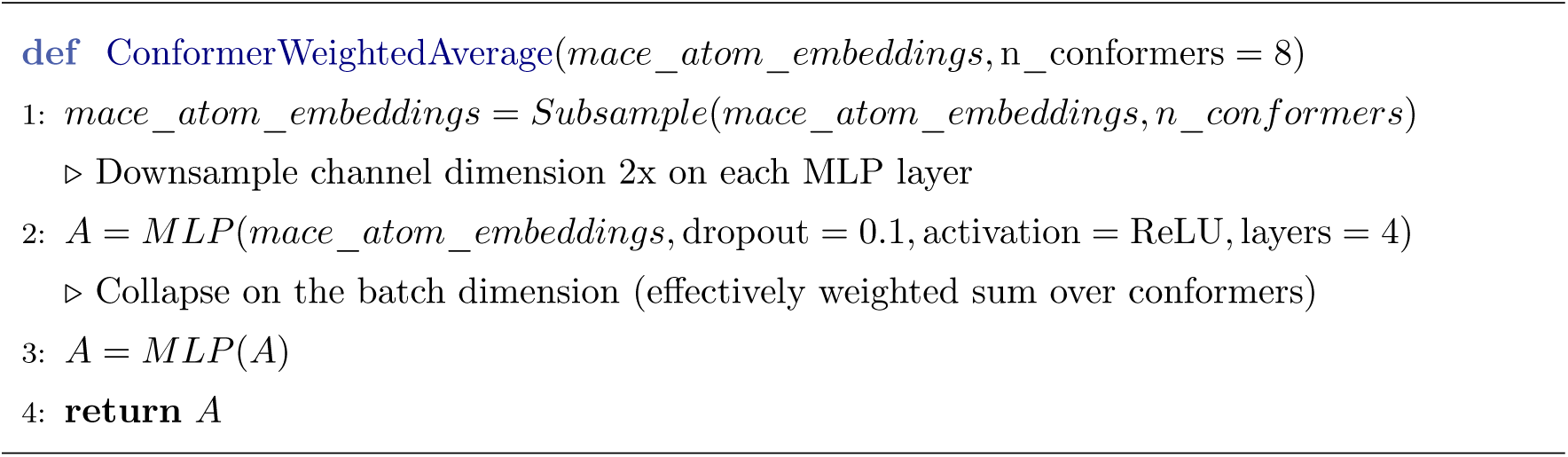

#### A.3.4 Confidence Head

We diverge from the published AF3 confidence head architecture. We mostly follow the confidence head implementation of [5] with some small modifications described below. We update the code to normalize the incoming logits using LayerNorms. We believe this implementation aligns closer to the implementation in the AF3 code release with minor changes in normalization strategy.

##### Algorithm 3 Confidence Head

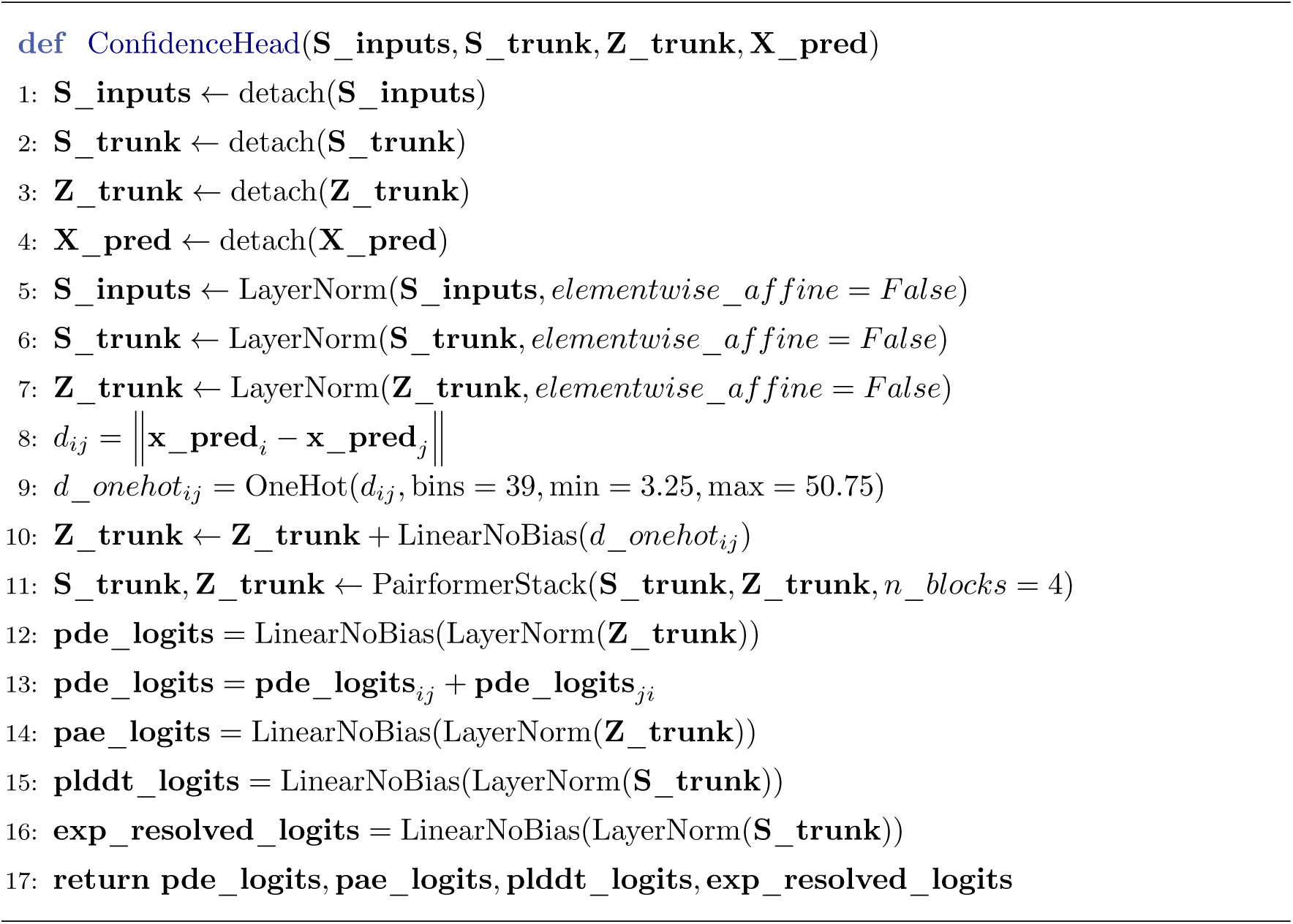

### A.4 Training Details

RF3 was trained in three separate stages: a) small crops using a 2021 date cut; b) large crops with a 2021 date cut; and c) large crops with a 2024 date cut. Comparing the model after step (b) with the final model lets us see the effect of 2.5 years of new structural data.

#### A.4.1 Training Dataset Mix

**Table S2:** Dataset sampling ratio mix.

| Dataset | Sampling Ratio |
| --- | --- |
| PDB (Chains/Interfaces) | 0.50 |
| AF2 Monomer Distillation | 0.34 |
| Multidomain Distillation | 0.10 |
| RNA Monomer Distillation | 0.02 |
| PDB (Disorder) | 0.02 |
| NA Complex Distillation | 0.02 |

In table S2, we detail the various datasets and corresponding sampling ratios used to train RF3.

For sampling from the Protein Data Bank (PDB), we adopt the weighted sampling scheme of [5] where we sample either from a dataset of chains or a dataset of interfaces, according to their relative weights. All other datasets are sampled uniformly. We do not explicitly show the inverse chirality dataset, since that is implemented as a train-time augmentation where we first sample from a dataset according to the probabilities above and then 2% of the time invert the chirality of all atoms in the structure.

#### A.4.2 Training Stages

The network was trained in three stages: first on crops of 384 tokens using the loss schema described above using the Adam optimizer with a learning rate of 1.8e-3 with 1000 warmup steps, then fine tune on crops of 768 tokens with a learning rate 9e-4 with no warmup and an additional loss for polymer-nonpolymer bonds, and finally with the same configuration as the second stage but with the date cutoff set to 1/2024. The first two stages only include structures deposited until 9/2021. The final results are all using an exponential moving average of the weights with a decay of 0.999.

The confidence head was trained with the main model frozen, using the exponential moving average weights of the main model. algorithm 3 shows the architecture of the confidence head. We used crop size 768 and learning rate 1.8e-3 with a 1000 step warmup. There was a separate confidence head training step for the 9/21 date cutoff model and the 1/24 model.

### A.5 Evaluation

For hyperparameter sweeps and assessing network performance during training, we made use of a validation dataset following [5] from that was disjoint from all evaluation sets. We then evaluated the model on three evaluation datasets.

All evaluations were performed with symmetry resolution analogous to that described in [5]. All methods were run with 10 recycles (adjusting from the default as needed), 5 diffusion samples per trunk output, and 200 diffusion steps.

#### A.5.1 Recent PDB Evaluation Set

For evaluation, we constructed a test dataset of recent PDB structures from after the latest training date cutoff of all models (1/2024). Broadly, we followed the procedure set forth in [5].

We begin by filtering PDB entries for quality; namely, we:

- Subset to PDB entries with release dates between 2024-01-01 and 2025-07-13
- Remove polymer chains with fewer than 4 resolved residues
- Remove entries with any clashing chains
- Remove entries that exceed 1,000 tokens, 10,000 atoms, or 4.5 Å in resolution

Next, we subset to low-homology chains and interfaces to evaluate. During evaluation, we predict all atoms within an entry, but only score chains and interfaces deemed sufficiently dissimilar from those seen during training. Following [5], we consider low homology to be <40% sequence identity for polymers and <0.85 Tanimoto similarity for ligands.

We then sample examples cluster-wise according to table S3. We find our test includes

**Table S3:**
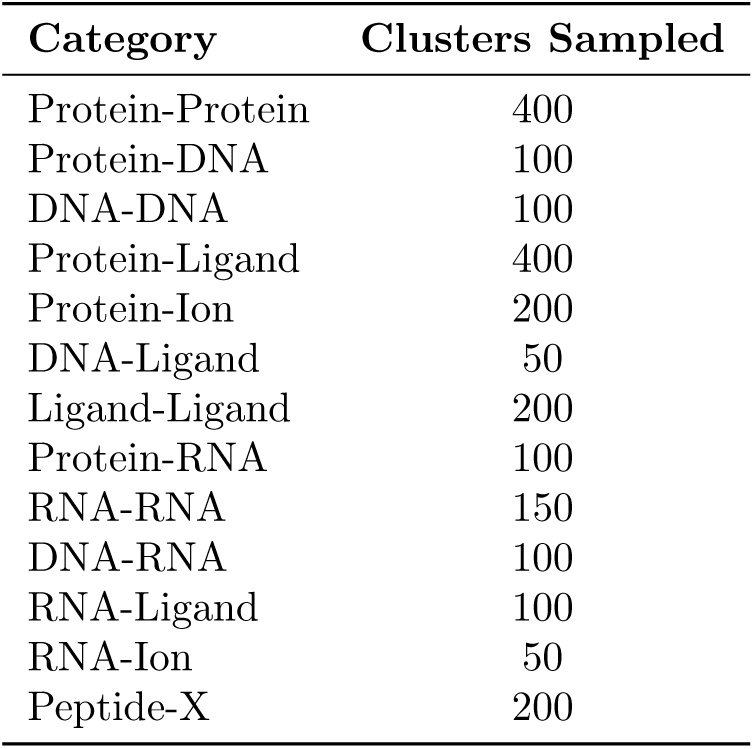
Number of interface clusters sampled for each category in the benchmark dataset.

#### A.5.2 Antibody Evaluation Set

The antibody evaluation set was constructed in an analogous manner to the Recent PDB Evaluation set, with the following additional steps:

1. We first used ANARCI [39] to search for antibody chains in all PDB entries represented in the structural antibody database (SAbDab) [40] as of July 27, 2025.
2. Next, we numbered these chains using the Chothia numbering system, and annotated the complementarity determining region (CDR) loops using the following inclusive bounds: LCDR1: [24, 34], LCDR2: [50, 56], LCDR3: [89, 97], HCDR1: [26, 32], HCDR2: [52, 56], HCDR3: [95, 102]. We defined antigens as any non-antibody proteins with a length of at least 30 that have at least one heavy atom within 5 Å of any CDR heavy atom.
3. These pairings were then used to subset the full set of interfaces identified from the PDB (following the definition of interfaces set forth in [5])
4. Last, we de-leaked the remaining interfaces against the training dataset by first sub-setting to examples released after the date cutoff (January 1, 2024) and then removing all interfaces in which the antigen shares a 40% sequence identity cluster with any training examples.

Note that this procedure is strictly more rigorous than the standard de-leaking procedure for interfaces (described above for the Recent PDB Evaluation Set) in which it is sufficient for either member of the interface to be de-leaked against the training data. Benchmarking results against this dataset shown in fig. S3

#### A.5.3 Mixed Chirality L/D Macrocyclic Peptide Evaluation Set

To evaluate performance of all models on a dataset of mixed-chirality peptides, we include the 11 examples deposited in the Cell 2022 paper, *Accurate de novo design of membrane-traversing macrocycles* [22]. Specifically, we include PDB entries: `7ubc, 7ubd, 7ube, 7ubf, 7ubg, 7ubh, 7ubi, 7uzl, 8cto, 8cun, 8cwa`. For evaluation, we ensure that these examples were not included in the training set of all models (e.g., we cannot evaluate Boltz-2 or RF3 trained through 1/2024 on this test set).

#### A.5.4 Input Preparation and Prediction

For each entry in the validation set, structural inputs were prepared starting from the corresponding `mmCIF` file obtained from the RCSB archive. Files were loaded via the AF3 from_mm-cif function with standard residue corrections (`fix_mse_residues=True, fix_arginines=True, fix_unknown_dna=True`), while excluding water molecules (`include_water=False`). Bond information was retained (`include_bonds=True`) and filtered to keep only those with type “covale” (covalent linkages), ensuring that other non-bonded components were excluded while retaining explicit covalent modifications such as post-translational modifications (PTMs). Crystallization-related ligands (as defined in the AF3 supplement) were excluded. The resulting structures were brought into the designated biological assembly using the `generate_bioassembly` function of AF3, with chain identities, assembly composition, and covalent bonds preserved. Pre-generated multiple sequence alignments were generated with MMSeqs2-GPU [33]; all methods used the same MSAs.

AF3 JSONs were produced from these assemblies and then converted into Boltz YAML and, for antibodies, Chai-1 FASTA, MSA parquet, and constraint files with a version of ABCFold [41] modified to preserve covalent constraints and assembly information. In particular, we adjust ABCFold by: mapping AF3 modifications to explicit CCD tokens (e.g., SEP, TPO, PTR), directly encoding ligand as CCDs, including multi-CCD glycans and ions, and collapsing identical chains into a single id: `[A,B,…]` block. We find such alternations are necessary to preserve covalent PTMs and ensure input consistency. For cyclic peptides, the `cyclic`: `true` flag was set in the Boltz YAML.

For Chai-1 FASTA conversion (antibody-only), our ABCFold AF3-Chai-1 modification enforces sequential A,B,C… chain IDs with while conserving the right bonded atom pair information, and reconstructs ligands, especially glycans, directly from bonded atom pairs into Chai-1 syntax. We also write Chai-native MSA parquet files from provided AF3 data json files.

All models were run on the full validation set under matched inference settings with 10 recycling steps and 5 diffusion samples ensuring identical recycling depth and batch processing to the other methods. All runs were executed on NVIDIA A100 GPUs, standardizing the computational environment. By preserving chain identities, assembly composition, covalent linkages, and MSA evidence across methods, we made every effort to ensure that differences in predictive performance reflected only model behavior, not discrepancies in input preparation or runtime configuration. All evaluations shown are computed after first applying symmetry resolution in accordance with [5] (benchmark scripts will be made available alongside the public code release).

### A.6 Training ProteinMPNN and LigandMPNN with AtomWorks

We copy the architecture code from ProteinMPNN [17] and LigandMPNN [18] and train both models from scratch within the AtomWorks framework. We train both LigandMPNN and Protein-MPNN, separately, on one A100 GPU for threes days. Sequence recovery performance, baselined against the prior models, given in fig. S5

### A.7 Supplementary Figures

**Fig. S1:**
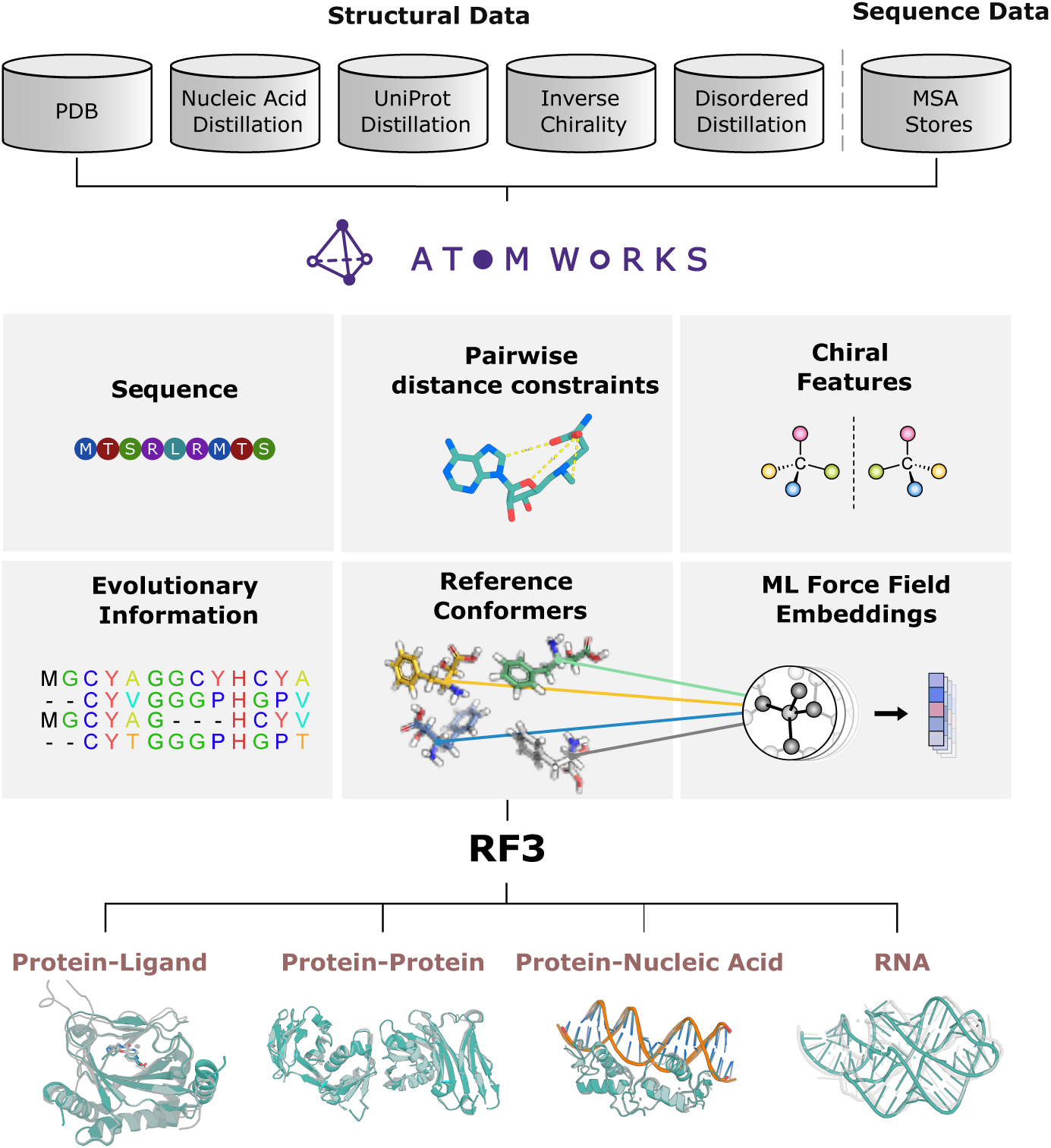
Datasets and features for RF3. **Top**: RF3 is trained on a diverse set of datasets including the Protein Data Bank (PDB), generated nucleic acid distillation sets, monomer distillation sets, PDB structures with inverted chirality, and PDB structures with extended disordered regions. **Middle**: The AtomWorks package parses all these disparate datasets and processes them through a single pipeline which can make a diverse set of features that are used for model training. **Bottom**: The model is trained to predict several biomolecular interactions including protein-ligand interactions, protein-protein interactions, protein-nucleic acid interactions, and RNA structure.

**Fig. S2:**
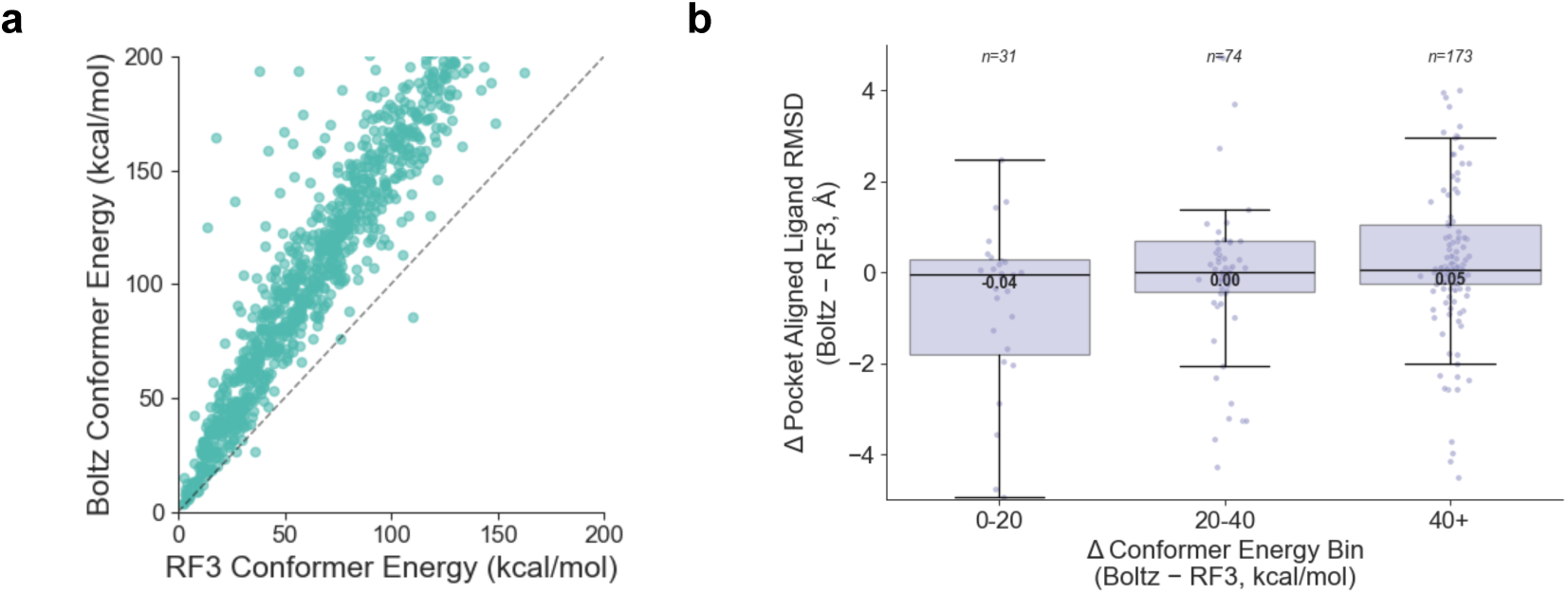
AtomWorks improves accuracy by using high-quality reference conformers. **A**. Comparison of reference conformer energies between the Boltz open source code and the RF3 method (reference conformer energies are computed using the Posebusters evaluation suite). **B**. Improvement of reference conformers correlates to accuracy. When subset to cases with fewer than 5 similar ligands in the PDB (by CCD code), we find that RF3 predictions are on average more accurate in cases where RF3 has lower reference conformer energies.

**Fig. S3:**
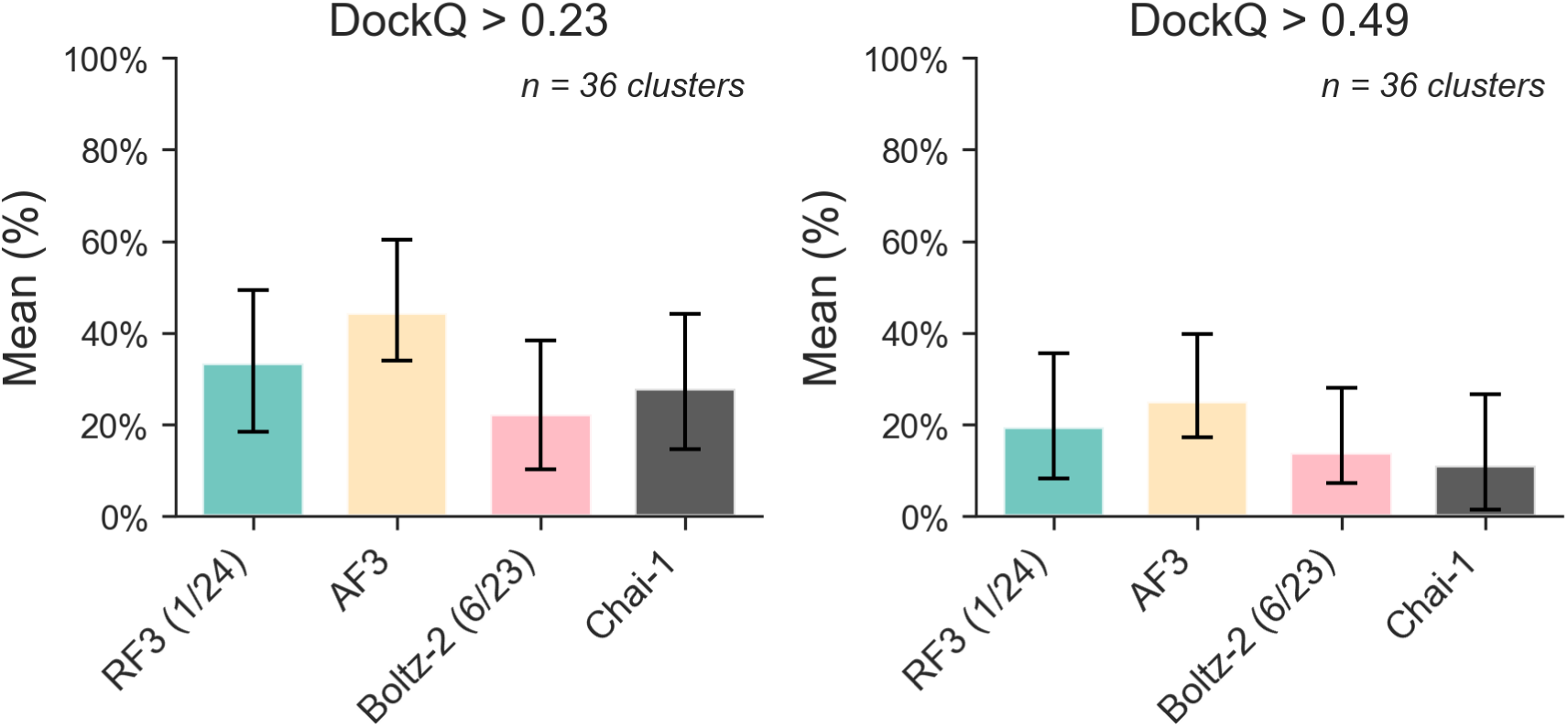
Evaluating structure prediction performance on antibody/antigen complexes. We predicted a test set of 75 unique PDB structures (58 after subsetting to those predicted and scored without error for all methods). We then clustered the predicted structures by antigen, using MMSeqs-2 with 40% sequence similarity threshold. After clustering, we retain 36 unique antigen clusters. DockQ scores are calculated for all examples; results reported are averaged within each cluster. We predict with a single model seed and five diffusion samples, choosing the model’s top-ranked structure by confidence score. All models are predicted with glycosylations. However, due to inconsistent treatment of leaving groups across models, we subset to only protein residues when computing DockQ scores. We make use of the PEPPR Biotite package [16] to compute DockQ.

**Fig. S4:**
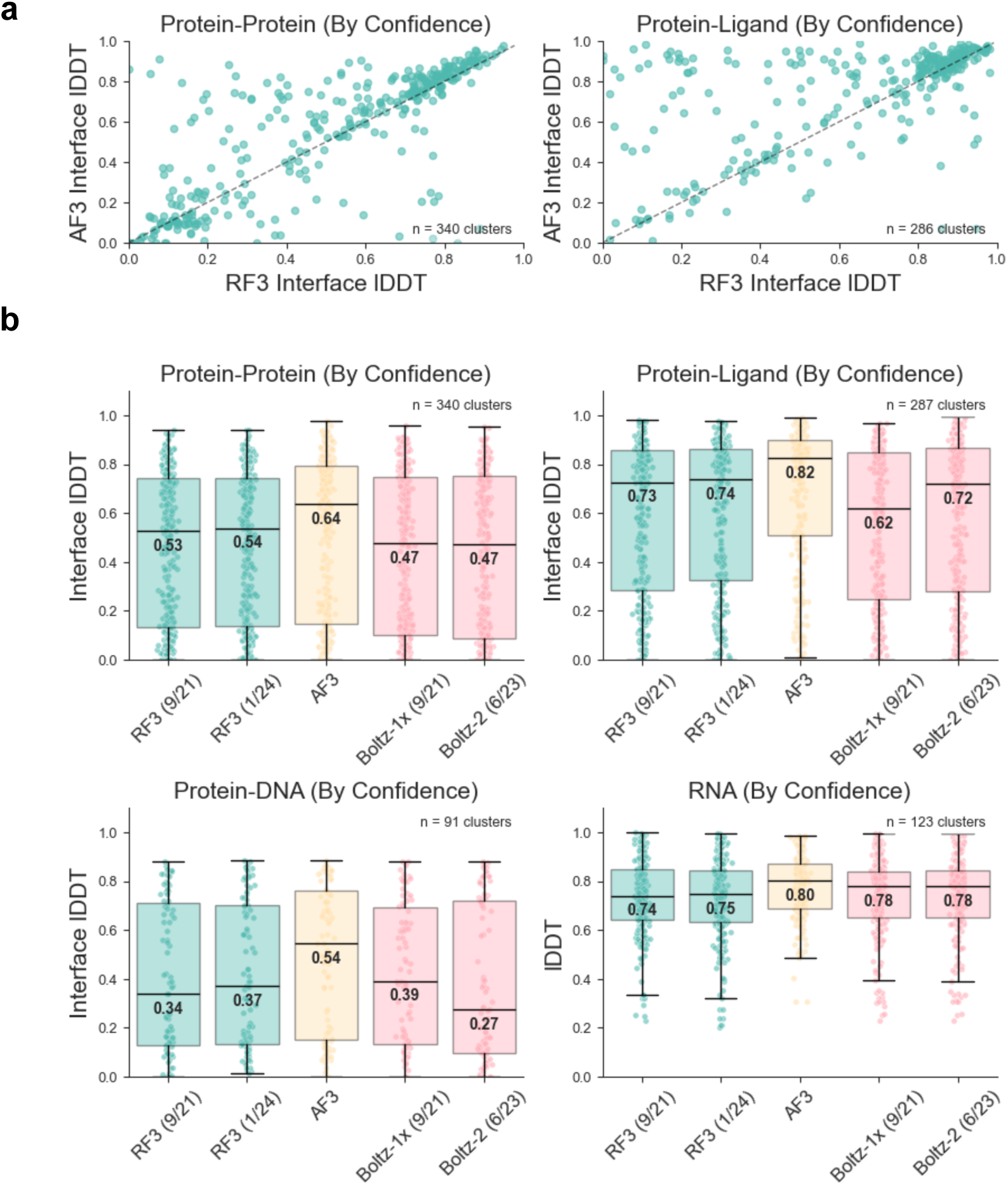
RF3 accurately predicts biomolecular interactions (selection by model confidence). **A**. Scatterplots comparing all-atom interface lDDT of RF3 and AF3 on protein-protein interactions and protein-ligand interactions. To reduce redundancy, the test dataset was clustered (by sequence homology 40% for polymers and CCD identity for non-polymers); each point represents a cluster mean. For all networks, we generate five structures from the same seed and use the model’s confidence head to select one structure for evaluation. **B**. Boxplots showing accuracy of RF3, AF3 and Boltz on different structure modeling tasks. Two versions of RF3 are shown: one trained on structures released before September, 2021 and another trained on structures deposited before January, 2024. Each point in the boxplot is a mean value over a cluster of structures. Model training date cutoff indicated in parentheses.

**Fig. S5:**
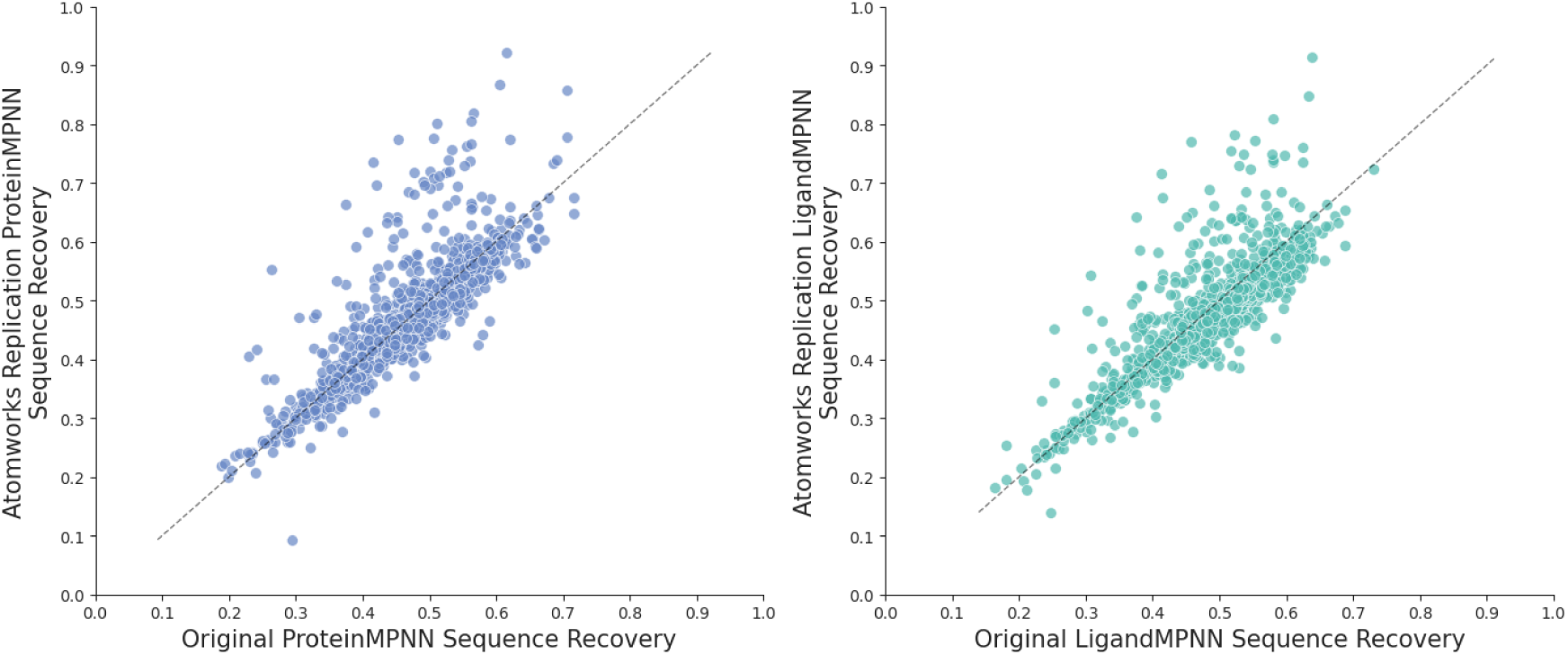
Comparison of original ProteinMPNN and LigandMPNN vs. AtomWorks replications. We find that naively integrating the ProteinMPNN and LigandMPNN architectures with RF3style dataloading and weighted sampling demonstrates comparable performance on a test set of recent low-homology sequences. We run 10 sequences per example (backbone) and calculate the sequence recovery per example; each point represents the mean across these ten examples.

**Fig. S6:**
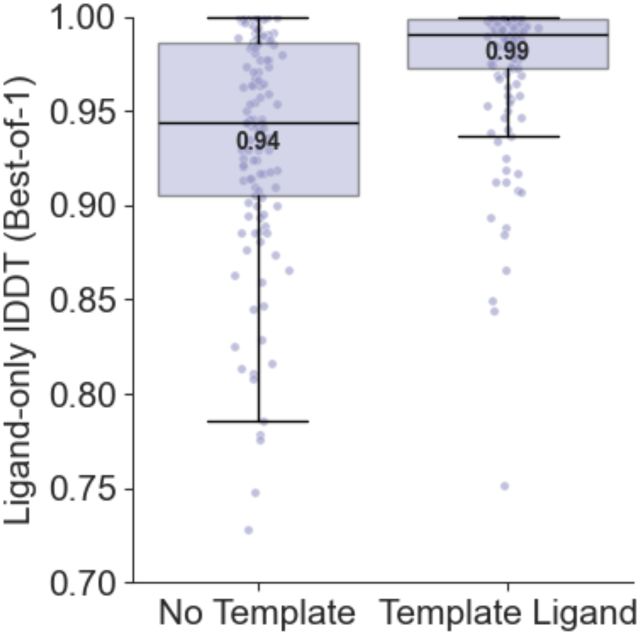
RF3 adheres to input ligand templates. On a set of 129 complexes in our test set, we find that providing ligand conformers increases ligand-only accuracy to 0.99 lDDT.

**Fig. S7:**
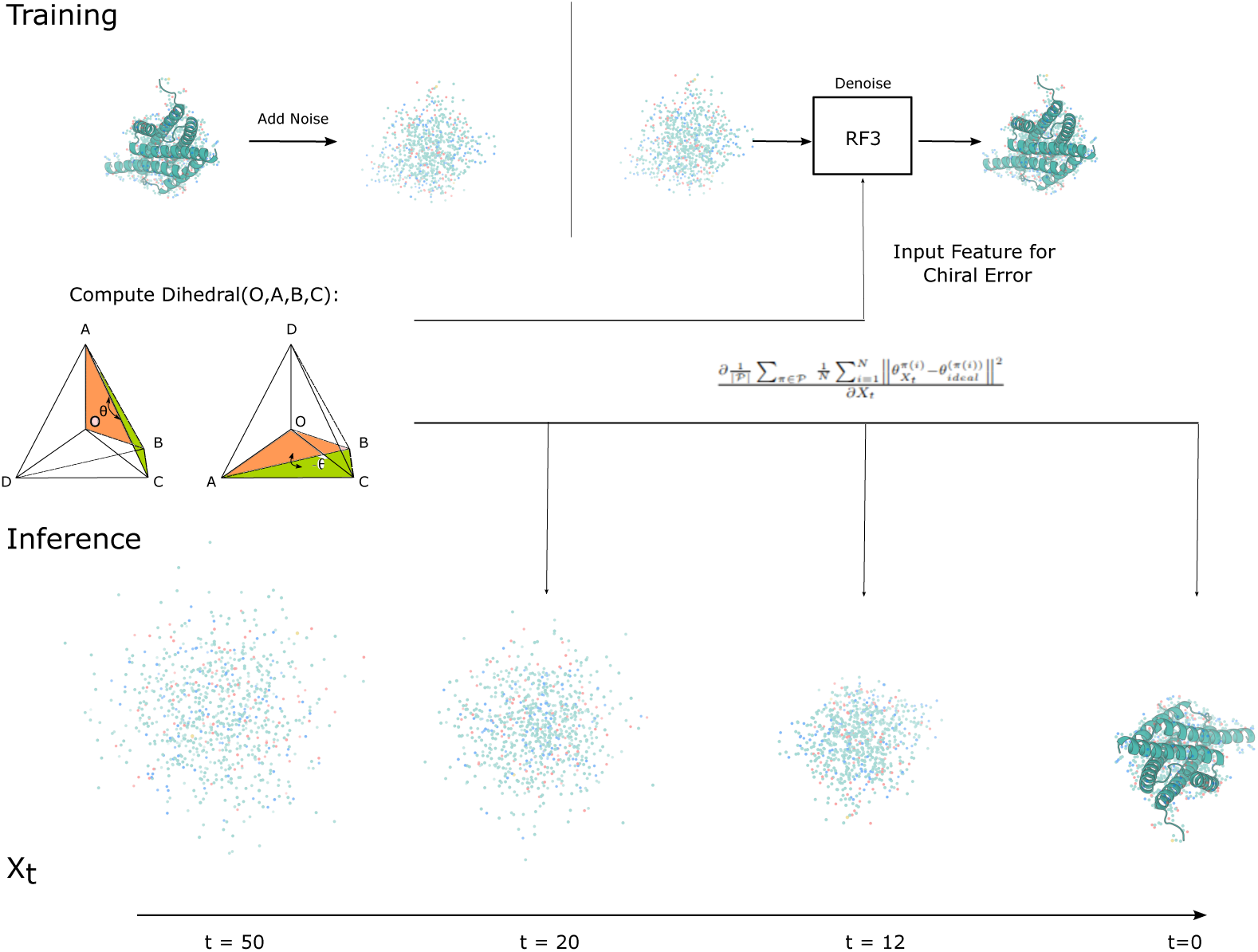
Depiction of chirality featurization in RF3. (Top) Procedure during training. A structure is sampled from the PDB and noise is added. The error of the angle of the tetrahedral chiral centers in the noisy input is provided as a separate feature to the network. (Bottom) Inference procedure. The structure is initialized as pure random gaussian noise and the network iteratively denoises the coordinates. At each denoising step, the error of the tetrahedral geometry is provided to the network in the same way it was provided in training.

**Fig. S8:**
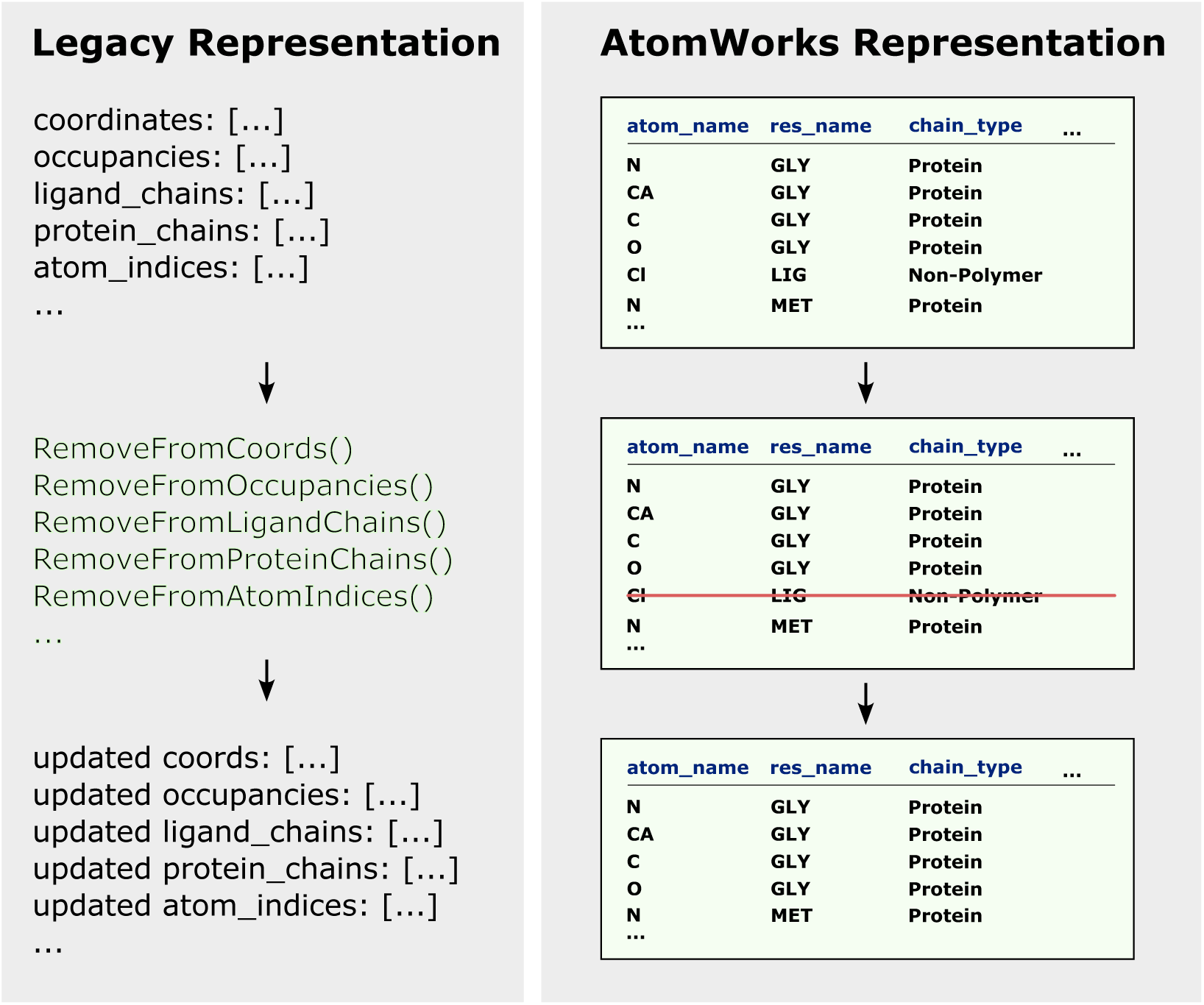
Comparison of new vs. old representations of structure for an illustrative step of removing a ligand atom (Cl). (left) Previously, pipelines were single functions that continuously converted input features into model tensors, discarding ground-truth information along the way. Researchers wishing to add new transforms (in this case, a transform that removes ligand atoms) had to grapple with complex features that might not be relevant to their tasks. (right) Within AtomWorks, our Transform-based approach reduces the complexity of implementing and modifying operations through a shared representation (the `AtomArray` from Biotite)

